# Radiative energy budgets of phototrophic surface-associated microbial communities and their photosynthetic efficiency under diffuse and collimated light

**DOI:** 10.1101/103705

**Authors:** Mads Lichtenberg, Kasper Elgetti Brodersen, Michael Kühl

## Abstract

We investigated the radiative energy budgets of a heterogeneous photosynthetic coral reef sediment and a compact uniform cyanobacterial biofilm on top of coastal sediment. By combining electrochemical, thermocouple and fiber-optic microsensor measurements of O_2_, temperature and light, we could calculate the proportion of the absorbed light energy that was either dissipated as heat or conserved by photosynthesis. We show, across a range of different incident light regimes, that such radiative energy budgets are highly dominated by heat dissipation constituting up to 99.5% of the absorbed light energy. Highest photosynthetic energy conservation efficiency was found in the coral sediment under light-limiting conditions and amounted to ~13% of the absorbed light energy. Additionally, the effect of light directionality, i.e., diffuse or collimated light, on energy conversion efficiency was tested on the two surface-associated systems. The effects of light directionality on the radiative energy budgets of these phototrophic communities were not unanimous but, resulted in local spatial differences in heat-transfer, gross photosynthesis and light distribution. The light acclimation index, E_k_ was >2 times higher in the coral sediment compared to the biofilm and changed the pattern of photosynthetic energy conservation under light-limiting conditions. At moderate to high incident 45 irradiances, the photosynthetic conservation of absorbed energy was highest in collimated light; a tendency that changed in the biofilm under sub-saturating incident irradiances, where higher photosynthetic efficiencies were observed under diffuse light. Our results suggest that the optical properties and the structural organization of phytoelements are important traits affecting the photosynthetic efficiency of biofilms and sediments.

## Introduction

Photosynthetic sediments and biofilms are characterized by pronounced vertical stratification of the microbial environment as a result of steep light gradients, high metabolic activity and limitations of heat and solute transport by diffusion (Kühl *et al*., 1996; Kühl and Fenchel, 2000; Al-Najjar *et al*., 2012). The radiative energy balance in such phototrophic microbial communities is affected by the incident radiative energy from the sun, of which a fraction is backscattered and thus not absorbed, while absorbed light energy is either photochemically conserved via photosynthesis or dissipated as heat via radiative energy transfer and non-photochemical quenching (Al-Najjar *et al*., 2010; Brodersen *et al*., 2014).

The quantity and quality of light are key regulating factors of photosynthesis, and the microscale distribution of light in microphytobenthic systems has been studied intensively over the last decades (Jørgensen and Des Marais, 1988; Lassen *et al*., 1992a; Kühl and Jørgensen, 1994; Kühl, 2005). A sub-saturating flux of photons will limit the rate of photosynthesis, as the available light is insufficient to support the maximal potential rate of the light reactions but as the photon flux increases the photosynthetic system saturates, whereby O_2_ becomes a competitive inhibitor on the binding-site of CO_2_ to Ribulose-1,5-bisphosphate carboxylase oxygenase (Rubisco) (Falkowski and Raven, 2007). In addition, when light energy absorption exceeds the capacity for light utilization, excess energy is channelled into heat production via non-photochemical quenching (NPQ) processes to avoid degradation of pigments and other cell constituents e.g. by reactive singlet oxygen produced by the de-excitation of triplet state chlorophyll (^3^Chl^*^) (Müller *et al*., 2001).

Photosynthetic organisms deploy different mechanisms to avoid photo damage, where NPQ is an effective short-term way to dispose of excess energy (Müller *et al*., 2001). If a photosynthetic cell experiences high light conditions on a daily basis, long-term regulation can be achieved by regulating the amount of light harvesting pigments (Nymark *et al*., 2009), where one strategy is to lower the light harvesting pigment content to reduce the absorption cross section by increasing transmittance, while another strategy involves upregulation of photoprotective pigments such as xanthophylls, that absorb energy-rich blue-green light but quench non-photochemically (Zhu *et al*., 2010).

Since photosynthetic cells perceive light from all directions, the angularity of light fields is important for determining the total radiance experienced by a cell (Kühl and Jørgensen, 1994), and it has e.g. been shown that the incident light geometry can influence photosynthetic light use efficiencies and photoinhibition in terrestrial plant canopies (Gu *et al*., 2002). In sediments, incident light will be spread by multiple scattering and, while the light field will become totally diffuse with depth (Kühl and Jørgensen, 1994), the response of benthic photosynthetic organisms to incident diffuse light is unknown. Through evaporation, an increase in cloud cover has been predicted with global warming (Schiermeier, 2006), which will potentially change the direction of light from relatively collimated beams (~85% in clear-sky conditions) to a more isotropic diffuse light field (~100% in cloud covered conditions)(Bird and Riordan, 1986; Brodersen *et al*., 2008; Gorton *et al*., 2010). In addition, submerged benthic systems will experience temporal and spatial differences in light field isotropy depending on turbidity, water depth, sun angle and the reflective properties of the surrounding environment (Brakel, 1979; Kirk, 1994; Wangpraseurt and Kühl, 2014).

Increased rates of photosynthesis have been observed in forest communities with an increasing proportion of diffuse light, possibly due to a more even distribution of light in the canopy (Gu *et al*., 1999; Krakauer and Randerson, 2003; Misson *et al*., 2005; Urban *et al*., 2007), whereby light energy is more efficiently harvested from all directions deeper in the canopy. This effect changes at the single leaf scale, where a 2-3% lower absorptance has been found under diffuse light as compared to collimated light at equivalent incident irradiances (Brodersen and Vogelmann, 2007). In corals, it has been observed that gross photosynthesis increased ~2-fold under collimated compared to diffuse light of identical downwelling irradiance (Wangpraseurt and Kühl, 2014) and the directional quality of light may thus elicit different photosynthetic responses and could potentially change the photosynthetic efficiency. A factor contributing to a possible difference in photosynthetic activity under diffuse and collimated light is photoinhibition, that initiates when the photosystems are saturated and the electron transport chain is fully reduced (Murata *et al*., 2007). Under high collimated light conditions, chloroplasts in leaves move to periclinal walls, and this might lead to reduced photoinhibition due to shading of other chloroplasts (Gorton *et al*., 1999). Under diffuse light, chloroplast movement to the periclinal walls is not complete (Williams *et al*., 2003) thus more randomly distributed, leading to possible less effective self-shading and photoprotection (Brodersen *et al*., 2008).

The balance between photosynthesis and respiration and as such, light use efficiency in benthic phototrophic systems is also influenced by the thickness of the diffusive and thermal boundary layers (Jørgensen and Des Marais, 1990; Jimenez *et al*., 2011; Brodersen *et al*., 2014). The diffusive boundary layer (DBL) is a thin water layer over a submerged object through which molecular diffusion is the dominant transport mechanism controlling the exchange of dissolved gases (e.g., O_2_ and CO_2_) and solutes with the ambient water (Jørgensen and Des Marais, 1990; Shashar *et al*., 1996) The DBL can thus impose a major control on respiration and photosynthesis in aquatic environments. Dissipation of absorbed solar radiation as heat drives an increase in surface temperature that is counter-balanced by heat transfer to the surrounding water via a thermal boundary layer (TBL), where convection dominates the transport of heat and the surface warming increases linearly with the incident irradiance (Jimenez *et al*., 2008). Heat and mass transfer phenomena through boundary layers are therefore important processes and parameters when considering rates of photosynthesis and radiative energy budgets.

In the present study, we present the first radiative energy budget of a heterogeneous coral reef sediment and compare it with the energy budget of a compact photosynthetic biofilm on a coastal sediment, and we investigate how diffuse and collimated light fields with identical levels of incident irradiance affect the radiative energy budget of the two microphytobenthic systems. Our analysis is based on a modified experimental approach first described by Al-Najjar *et al*. (2010).

## Materials and methods

### Sample sites and collection

Coral reef sediment was sampled in April 2012 from a sheltered pseudo-lagoon (‘Shark Bay’ (Werner *et al*., 2006) on the reef flat surrounding Heron Island (151°55’E, 23°26’S) that is located on the southern boundary of the Great Barrier Reef, Australia. Maximal incident solar photon irradiance at 130 the sediment surface of the shallow reef flat during calm mid-day low tides is ~1500-2000 µmol photons m^−2^ s^−1^ (Jimenez *et al*., 2012; Wangpraseurt *et al*., 2014b). The coral sediment (CS) was mainly composed of bright, semi-fine grained particles (mostly in the 200-500 µm size fraction) of deposited CaCO_3_ from decomposed corals and other calcifying reef organisms. Diatoms, dinoflagellates and cyanobacteria were found as aggregates in the sediment pore space along with amorphic organic material (detritus) throughout the upper few mm of the sediment (Fig. S1).

The biofilm (BF) originated from a shallow sand bar at Aggersund, Limfjorden (Denmark) experiencing maximum summer photon irradiance of 1000-1500 µmol photons m^−2^ s^−1^. The biofilm was comprised of a ~1 mm thick smooth layer of photosynthetically active cyanobacteria and microalgae embedded in exopolymers on top of fine-grained (125-250 µm) dark sulfidic sandy sediment (Nielsen *et al*., 2015). The porosity of the coral sediment and biofilm, ϕ, was 0.78 and 0.80, respectively, as determined from the weight loss of wet sediment (known initial volume and weight) after drying at 60°C until a constant weight was reached:

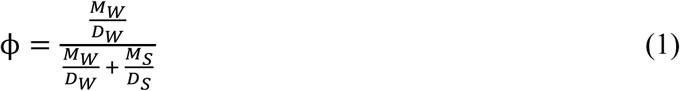

where M_W_ is the weight of water, D_W_ is the density of water, M_S_ is the weight of sediment/biofilm, and D_S_ is the sediment/biofilm density.

### Coral Sediment samples

The CS samples were collected with Perspex corers (inner diameter 5.3 cm), and were maintained under a continuous flow of aerated seawater at ambient temperature and salinity (26°C and S=35) under a natural solar light regime for ~24 h prior to further handling at the Heron Island Research Station (HIRS), Australia. Sediment cores were then mounted in a custom-made flow-chamber flushed with aerated seawater (26°C and S=35) for another 24 h prior to measurements. During the acclimation time in the flow-chamber, the sediment cores were kept under a downwelling photon irradiance of ~1000 µmol photons m^−2^ s^−1^ provided by a fiber-optic tungsten halogen lamp equipped with a collimating lens (KL2500-LCD, Schott GmbH, Germany). Before measurement at each experimental irradiance, the coral sediment core was illuminated for at least 45 minutes to ensure steady state O_2_ and temperature conditions; as confirmed from repeated microprofile measurements. Throughout measurements, the flow-chamber was flushed with a stable laminar flow (~0.5 cm s^−1^) of filtered aerated seawater over the sediment surface as generated by a Fluval U1 pump submerged in a 20L thermostated aquarium (26°C and S=35) and connected with tubing to the flow-chamber.

### Biofilm samples

The BF samples were collected and contained in small rectangular plastic trays (7 ⨯ 2 ⨯ 5 cm) with the upper surface exposed and flush with the upper edge of the tray wall. After collection, the samples were kept humid and under a 12:12 h light-dark regime (~100 µmol photons m^−2^ s^−1^) in a thermostated room (16-18°C). The biofilm surface appeared dark green–brownish due to predominance of dense communities of cyanobacteria and diatoms (Lassen *et al*., 1992b). Prior to measurements, a sample tray was placed for 2 days in a flow-chamber flushed with 0.2 µm filtered aerated seawater (21°C, S=30) under a downwelling photon irradiance of ~500 µmol photons m^−2^ s^−1^. During measurements, a stable laminar flow (~0.5 cm s^−1^) over the biofilm surface was maintained by a water pump (Fluval U1, Hagen GmbH, Germany) immersed in an aquarium with filtered aerated seawater (21°C, S=30) and connected with tubing to the flow-chamber.

### Experimental setup

Illumination was provided by a fiber-optic tungsten halogen lamp equipped with a collimating lens (KL-2500 LCD, Schott, Germany) positioned vertically above the flow-chamber. The intensity of the lamp could be regulated without spectral distortion by a built-in filter wheel with pinholes of various sizes. The downwelling photon irradiance of photosynthetically active radiation (PAR, 400-700nm), Ed(PAR), (see definitions of abbreviations) was measured with a calibrated irradiance meter (ULM-500, Walz GmbH, Germany) equipped with a cosine collector (LI-192S, LiCor, USA). Defined experimental irradiances (0, 50, 100, 200, 500 and 1000 µmol photons m^−2^ s^−1^) were achieved by adjusting the aperture on the fiber-optic lamp. The downwelling spectral irradiance at the above-mentioned levels was also measured in radiometric energy units (in W m^−2^ nm^−1^) with a calibrated spectroradiometer (Jaz, Ocean Optics, USA).

Collimated light was achieved by attaching a collimating lens to the fiber cable of the lamp. Diffuse light was achieved by inserting a TRIMMS diffuser (Transparent Refractive Index Matched Microparticles) (Smith *et al*., 2003) between the collimator and the sample followed by lamp adjustment to achieve the same absolute levels of downwelling irradiance on the biofilm/sediment surface in collimated and diffuse light treatments.

### Microscale measurements of O_2_ and temperature

*Oxygen concentrations* were measured with a Clark-type O_2_ microsensor (tip diameter ~25 µm, OX-25, Unisense AS, Aarhus, Denmark) with a fast response time (<0.5 s) and a low stirring sensitivity (<1-2%) (Revsbech, 1989). The microsensor was connected to a pA-meter (Unisense A/S, Aarhus, Denmark) and was linearly calibrated at experimental temperature and salinity from measurements in the aerated seawater in the free-flowing part of the flow-chamber and in anoxic layers of the sediment.

Temperature measurements were performed with a thermocouple microsensor (tip diameter ~50 µm; T50, Unisense A/S, Aarhus, Denmark) connected to a thermocouple meter (Unisense A/S, Aarhus, Denmark). The temperature microsensors were linearly calibrated against readings of a high precision thermometer (Testo 110, Testo AG, Germany; accuracy ±0.2°C) in seawater at different temperatures. Analogue outputs from the temperature and O_2_ microsensor meters were connected to an A/D converter (DCR-16, Pyroscience GmbH, Germany), which was connected to a PC. All microsensors were mounted in a PC-interfaced motorized micromanipulator (MU-1, PyroScience, GmbH, Germany) controlled by dedicated data acquisition and positioning software (ProFix, Pyroscience, Germany). The micromanipulator was oriented in a 45° angle relative to the vertically incident light to avoid self-shading, especially in the light measurements. Depth profiles of temperature and O_2_ concentration were measured in vertical steps of 100 µm. Before profiling, the microsensor tips were manually positioned on the sample surface to define the z=0 position, while observing the biofilm/sediment surface through a stereo dissection microscope.

The local volumetric rates of gross photosynthesis (P_G_(z); in units of nmol O_2_ cm^−3^ s^−1^) were measured with O_2_ microsensors using the light-dark shift method (Revsbech and Jørgensen, 1983). Volumetric rates were measured in vertical steps of 100 µm throughout the sediment until no photosynthetic activity in the given depth was detected. The immediate O_2_ depletion rate upon brief (2-4 s) darkening equalled the local rate of photosynthesis just prior to darkening; while no response in the O_2_ signal upon darkening indicated a zero rate of photosynthesis. Areal rates of gross photosynthesis (in nmol O_2_ cm^−2^ s^−1^) were calculated by depth integration over the euphotic zone with respect to the measuring interval used in the depth profile measurement of P_G_(z), similar to Al-Najjar *et al*. (2010); Al-Najjar *et al*. (2012):

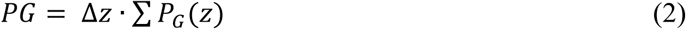

### Temperature and O_2_ calculation

The net upward flux of O_2_ from the photic zone of the sediments into the overlaying seawater was calculated (in nmol O_2_ cm^−2^ s^−1^) from measured steady-state O_2_ concentration profiles using Fick’s first law of diffusion:

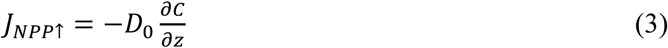

where D_0_ is the diffusion coefficient of O_2_ in seawater at experimental temperature and salinity and 
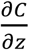
 is the linear O_2_ concentration gradient in the DBL.

The downward O_2_ flux from the photic zone of the sediments to the aphotic part of the sediment/biofilm was calculated in a similar manner as:

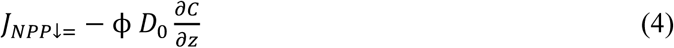

The total flux of O_2_ out of the photic zone, i.e., the total net photosynthesis in the photic zone (NPP), was subsequently calculated as the difference between the upward and downward O_2_ flux (Kühl *et al*., 1996).

To calculate the radiative energy conserved via photosynthesis (J_PS_) (in J cm^−2^ s^−1^) we multiplied the areal gross photosynthesis, GPP, with the Gibbs free energy formed in the light-dependent reactions, where O_2_ is formed by splitting water, which gains (including the formation of ATP) a Gibbs free energy of E_G_ = 482,9 kJ (mol O_2_)^−1^ (Thauer *et al*., 1977).

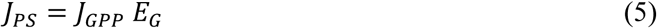

The amount of the absorbed light energy that was not photochemically conserved was dissipated as heat resulting in a local increase of the sediment/biofilm temperature relatively to the ambient seawater and thereby leading to the establishment of a thermal boundary layer (TBL). The heat dissipation, i.e., the heat flux (in J m^−2^ s^−1^) from the sediment/biofilm into the water column was calculated by Fourier’s law of conduction:

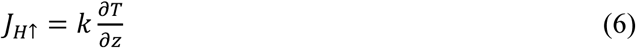

where *k* is the thermal conductivity in seawater (0.6 W m^−1^ K^−1^) and *dT/dz* is the measured linear temperature gradient in the TBL (Jimenez *et al*., 2008). The heat flux from the photic zone into the aphotic sediment/biofilm, J_H↓_., was calculated as in Eq. 6 but with the thermal conductivity constant of the sediment, *k(b)*, which was estimated as:

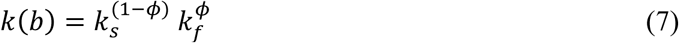

where *k_s_* is the carbonate thermal conductivity (3.1 W m^−1^ K^−1^;(Clauser and Huenges, 1995)), *k_f_* is the seawater thermal conductivity, and ϕ is the porosity of the sediment (Lovell, 1985).

The total heat flux, was used as an estimate of the total heat dissipation in the photic zone and was calculated as: ∑*J*_*H*_ = *J*_*H*↑_ − *J*_*H*↓_

### Microscale light measurements

Spectral photon scalar irradiance was measured in units of counts nm^−1^ with a fiber-optic scalar irradiance microprobe (integrating sphere diameter ~100 µm; (Lassen *et al*., 1992a)) connected to a fiber-optic spectrometer (USB2000, Ocean Optics, Dunedin, FL, USA). A black non-reflective light-well was used to record spectra of the downwelling photon scalar irradiance, *E_d_(λ)*, (in units of counts nm^−1^) with the tip of the scalar irradiance microsensor positioned in the light path at the same distance from the light source as the sediment surface. Using identical light settings, the absolute downwelling irradiance, *E_ABS_(λ)* (in W m^−2^) was also quantified with a calibrated spectroradiometer (Jaz-ULM, Ocean Optics, Dunedin, Florida, USA).

### Irradiance calculations

The spectral scalar irradiance, *E_0(λ)_*, was measured in vertical steps of 0.1-0.2 mm in the sediment and was calculated as the fraction of the incident downwelling irradiance, i.e., *E_0_(λ)/E_d_(λ)*, and plotted as transmittance spectra in % of *E_d_(λ)*. The relative measurements of scalar irradiance in different depths in the biofilm/sediment were converted to absolute scalar irradiance spectra in units of W m^−2^ nm^−1^ as *E_ABS_(λ)*E_0_(λ)/E_d_(λ)*. Absolute scalar irradiance spectra were converted to photon scalar irradiance spectra (in units of µmol photons m^−2^ s^−1^ nm^−1^) by using Planck’s equation:

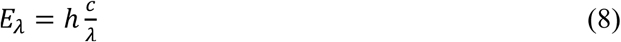

Where *E_λ_* is the energy of a photon with wavelength, *λ, h* is Planck’s constant (6.626 × 10^−34^ W s^2^), and *c* is the speed of light in vacuum (in m s^−1^).

Spectral attenuation coefficients of scalar irradiance, K_0_(λ), were calculated as (Kühl, 2005):

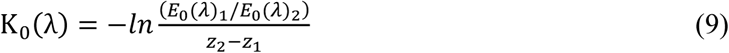

where *E_0_(λ)_1_* and *E_0_(λ)_2_* are the spectral scalar irradiances measured at depth z_1_ and z_2_, respectively. Light attenuation was also calculated by integrating the spectral quantum irradiance over PAR (420-700 nm) yielding the PAR scalar irradiance (*E_0_(PAR)*), in µmol photons m^−2^ s^−1^), i.e., the light energy available for oxygenic photosynthesis at each measurement depth. The diffuse attenuation coefficient of *E_0_(PAR)*, *K_0_(PAR)*, was obtained by fitting the measured *E_0_(PAR)* vs. depth profiles with an exponential model:

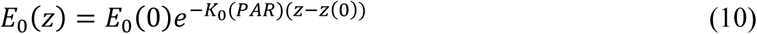

### Reflectance measurements

The PAR irradiance reflectance (R) of the sediment/biofilm surface was calculated as

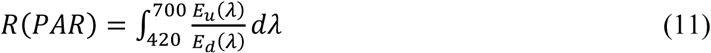

where *E_u_(λ)* is the upwelling irradiance at the sediment surface, here estimated as the diffuse backscattered spectral radiance measured at the sediment surface (Kühl, 2005) and *E_d_(λ)* is the downwelling irradiance estimated as the backscattered spectral radiance measured over a white reflectance standard (Spectralon; Labsphere, North Sutton, NH, USA); both measured with a fiber optic field radiance microprobe (Jørgensen and Des Marais, 1988). The R(PAR) measurements assumed that the light backscattered from the sediment/biofilm surface was completely diffused (Kühl and Jørgensen, 1994).

### Absorbed light energy

The absorbed light energy (JABS)(in W m^−2^ = J m^−2^ s^−1^) in the sediment/biofilm was estimated by subtracting the downwelling and upwelling irradiance at the surface:

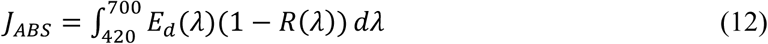

where *E_d_(λ)* and *R(λ)* are the downwelling spectral irradiance and irradiance reflectance, respectively. This parameter is equivalent to the so-called vector irradiance, which is a measure of the net downwelling radiative energy flux.

### Energy budget and photosynthetic efficiency calculations

A balanced radiative energy budget of the sediment/biofilm was calculated according to (Al-Najjar *et al*., 2010) with slight modifications (Fig. 1) as:

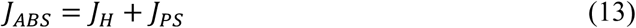

assuming that autofluorescence from the sediment/biofilm was negligible. Consequently,*ɛ_PS_*+*ɛ_PS_*=1, where *ɛ_PS_* and *ɛ_H_* represent the efficiency of photosynthetic energy conservation and heat dissipation, respectively, for a given absorbed light energy JABS in the entire euphotic zone (Al-Najjar *et al*., 2010):

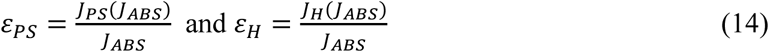

**Figure 1.**
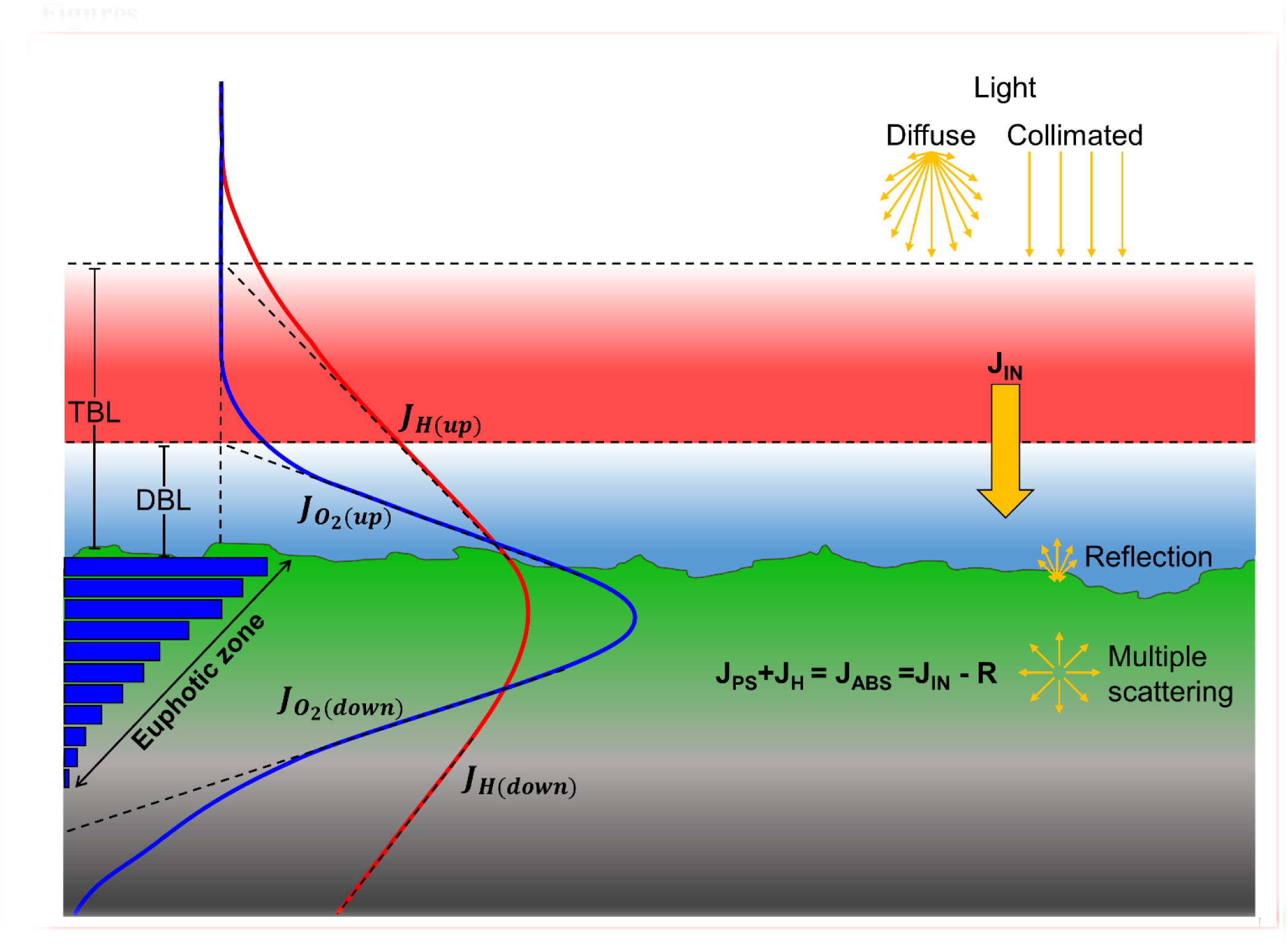
**Major pathways of light energy conversion and dissipation in biofilm and coral sediment.** Incident irradiance was either diffuse or collimated (top yellow arrows) and supplied the sediments with the incoming energy flux, J_IN_ (solid yellow arrow). A fraction of the incoming light energy was reflected from the surface and thereby not a part of the absorbed light energy (J_ABS_). Through multiple scattering by biotic and abiotic components in the biofilm/sediment, the light field becomes increasingly diffuse with depth. The absorbed light energy is either photochemically conserved in photosynthesis (J_PS_) in the photic zone or dissipated as heat (J_H_) via radiative energy transfer and non-photochemical quenching leading to local heating in the biofilm/sediment (red line). Gross photosynthesis (blue bars) is dependent on light and is thus higher near the surface which drives a production of O_2_ (blue line) that exceeds the consumption via respiration and leads to the formation of a diffusive boundary layer (DBL). The surplus of O_2_, i.e., the net photosynthesis, can be calculated as the sum of the upwards (*J_o2(Up)_*) and downwards (*J_o2(Down)_*) flux of O_2_ from the photic zone. Similarly, the fraction of the absorbed energy that is dissipated as heat can be calculated as the sum of upwards (*J_H(Up)_*) and downwards (*J_H(Down)_*) heat flux through the thermal boundary layer (TBL) into the overlaying water and into the aphotic sediment/biofilm layer, respectively.

Areal gross photosynthesis rates as a function of *J_ABS_*, were fitted with the saturated exponential model (Webb *et al*., 1974) to estimate the maximum conserved energy flux by photosynthesis (*J_PS, max_*)(in J m^−2^ s^−1^):

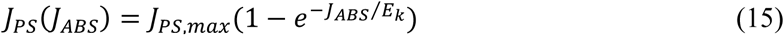

This yielded an estimate of the maximum photochemically conserved energy flux *J_PS, max_*. The respective efficiencies under light-limiting conditions, i.e., for *J_ABS_*→0, were then calculated as:

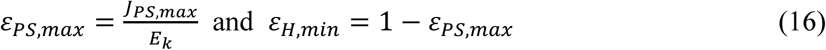

where *E_k_* is the photochemical light acclimation index, i.e., the irradiance at the onset of photosynthetic saturation, calculated as *E_k_*=*J_PS, max_*/α, where α is the initial slope of the fitted photosynthesis vs. *J_ABS_* curve.

## Results

### Light measurements

At all incident irradiances, the photon scalar irradiance, E0(PAR), was enhanced to 120-200% immediately below the surface (0-0.2 mm in the biofilm; 0-1 mm in the coral sediment) relative to the incident downwelling photon irradiance (Fig. 2). Light attenuation was strongly enhanced around wavelengths 625 nm and 670 nm, corresponding to absorption maxima of phycocyanin and Chl a (Fig. 3). Surface reflection from the biofilm surface was on average 1.8% and 1.7% of the incident PAR under diffuse and collimated light, respectively, while it was >15 times higher in the coral sediment, i.e., 30.2% and 28.1% for diffuse and collimated light, respectively. Reflection did not change with increasing irradiance (Fig. S2).

**Figure 2.**
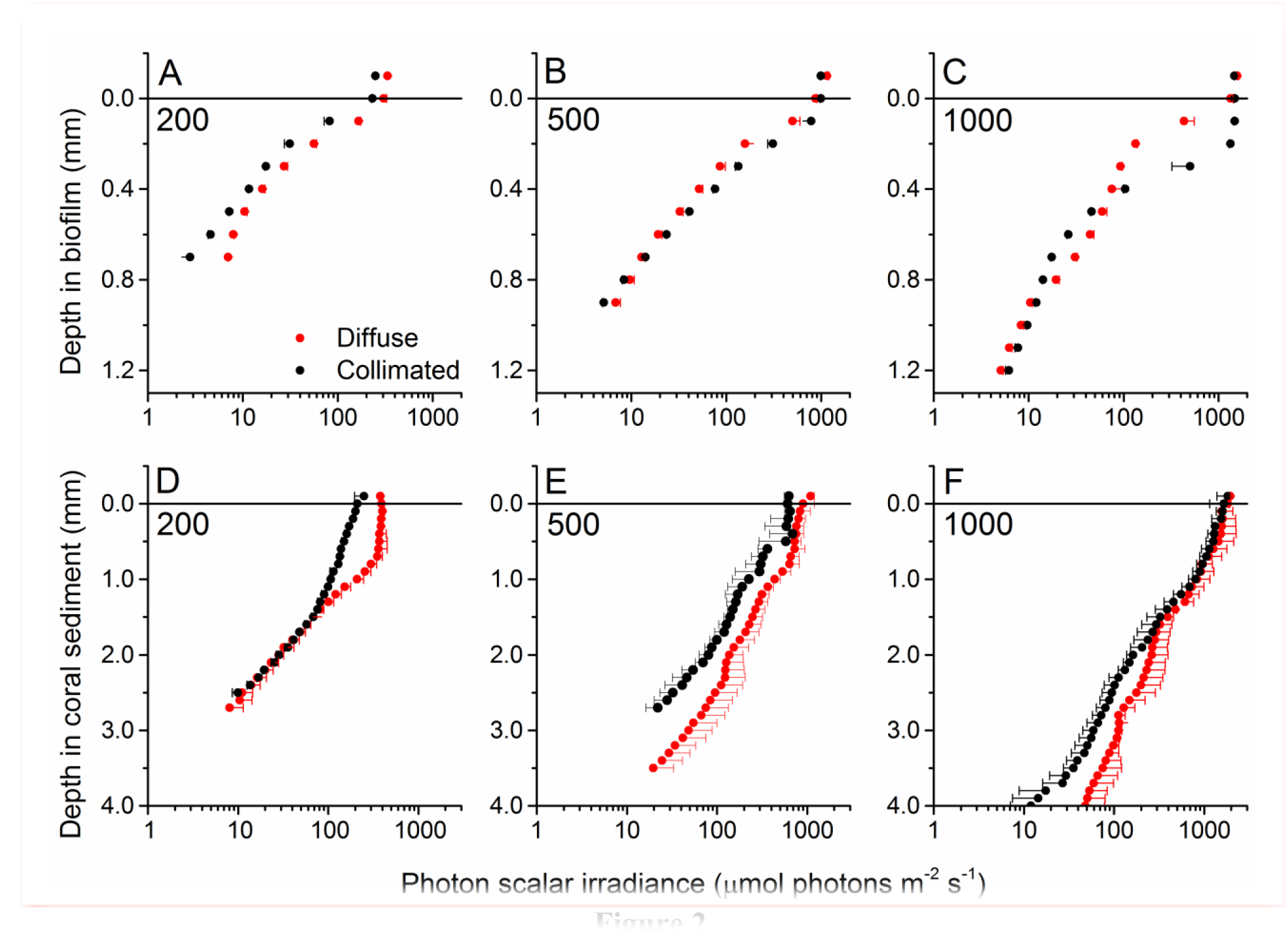
**Vertical profiles of photon scalar irradiance** (PAR, 400-700 nm) in the biofilm (A-C) and coral sediment (D-F) under different incident photon irradiance (numbers in panels) of collimated (black symbols) and diffuse (red symbols) light. In the biofilm the light attenuation coefficient (α) was estimated in both the upper (0-0.4mm) and lower (0.5-1.2mm) layer while. was estimated for the entire exponential part of the curve in the coral sediment (R^2^>0.95 in all cases). Data points show averages ± 1 S. D. (note that for clarity, only plus S. D. is shown for diffuse light and minus S. D. for collimated light; *n*=3).

**Figure 3.**
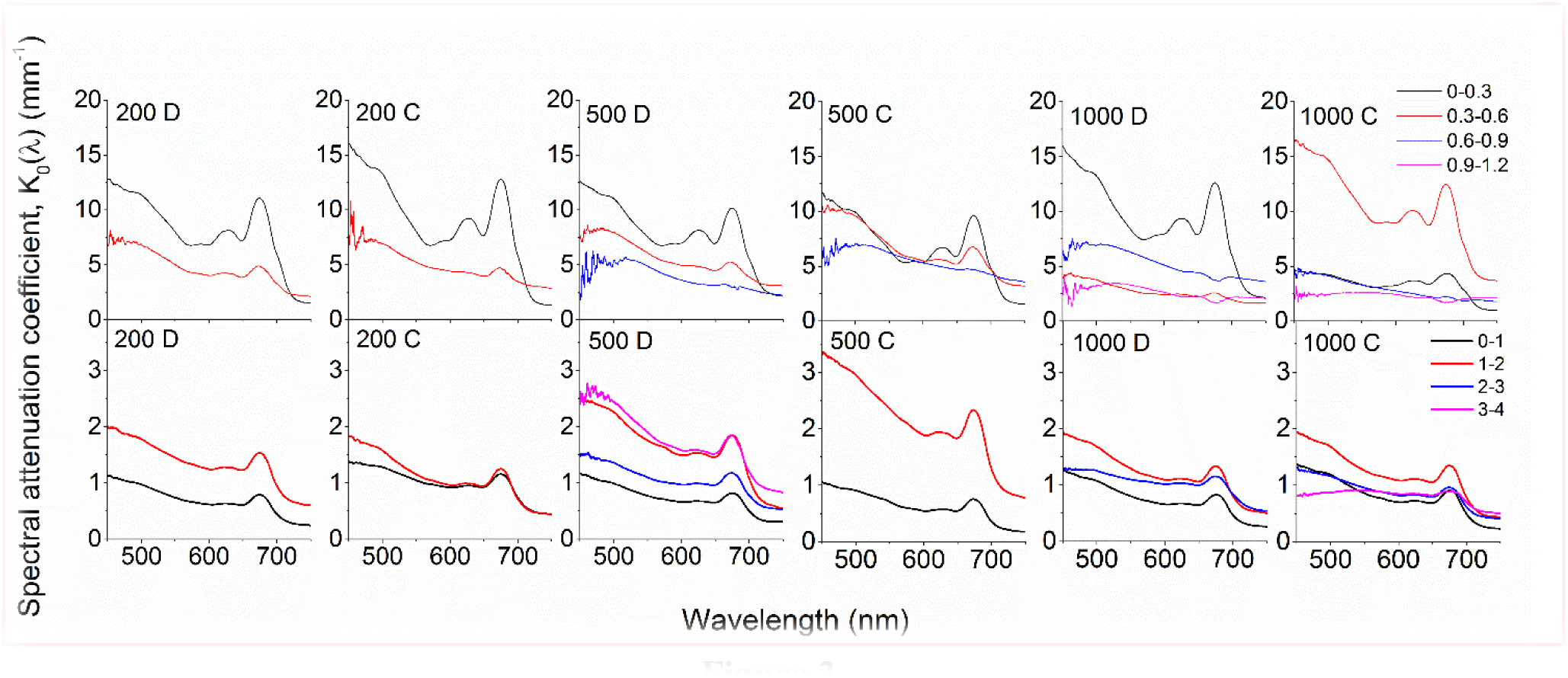
**Spectral attenuation coefficients,** K0(λ)(PAR) of photon scalar irradiance calculated over 300 µm (biofilm; upper panels) and 1000 µm (coral sediment; lower panels) depth intervals. Numbers in panels indicate incident photon irradiance in µmol photons m^−2^ s^−1^, while the letters C and D denote collimated and diffuse incident light, respectively. Curves represent averages (*n=3; S.D. not shown for clarity*).

Below the uppermost sediment and biofilm layers exhibiting local photon scalar irradiance enhancement, light was attenuated exponentially with depth (Fig. 2). At the highest incident photon irradiances (500 and 1000 µmol photons m^−2^ s^−1^), the exponential attenuation of collimated light within the biofilm was observed below 0.2 mm, whereas diffuse light was attenuated exponentially from the biofilm surface under all investigated irradiance levels (Fig. 2). In the coral sediment, the exponential attenuation occurred deeper (below 0.5-0.7 mm) due to enhanced scattering, redistribution and trapping of photons in the upper sediment layers (Fig. 2). In the biofilm, PAR attenuation was stronger in the top layer than in the bottom layer both for diffuse and collimated light (Fig. 2). Additionally, attenuation of collimated light in the top layer was stronger than for diffuse light at all irradiances except 1000 µmol photons m^−2^ s^−1^, whereas light attenuation in the lower sediment dominated layers was similar for diffuse and collimated incident light. In the coral sediment no distinct differences in light attenuation was observed between top-and bottom layers other than a deeper onset of exponential attenuation (0.5-0.7 mm). The top layer of the biofilm showed ~10 times stronger light attenuation than the coral sediment with average PAR attenuation coefficient of α= 9.52 mm^−1^ and α= 10.54 mm^−1^ for diffuse and collimated light, respectively, compared to α= 1.18 mm^−1^ in the coral sediment (both light types).

In both sediments, attenuation of light corresponded to absorption maxima of Chl a (440 and 670 nm) and phycocyanin (620 nm) (Fig. 3). A third attenuation maximum was observed around 575 nm indicative of phycoerythrin, commonly found in cyanobacteria (Colyer *et al*., 2005). In the biofilm, attenuation of visible light was strongest in the top 0.3 mm of the biofilm, except under the highest collimated irradiance (1000 µmol photons m^−2^ s^−1^), where the strongest attenuation occurred over the 0.3-0.6mm zone (Fig. 3). Below 0.6 mm, the enhanced attenuation around wavelengths 575 nm, 625 nm and 670 nm decreased and the attenuation of light in the PAR region became more uniform in the underlying layers (Fig. 3). Again, attenuation of collimated light was slightly higher than diffuse light. In the coral sediment, the highest light attenuation was 1-2 mm below the sediment surface (~1.6 mm^-1^ at 670 nm at all incident irradiances) while the lowest attenuation was found in the upper 0-1mm, consistent with the scalar irradiance profiles (Fig. 2 and 3).

### Temperature and O_2_ microenvironment

In the biofilm, a ~0.8 mm thick diffusive boundary layer (DBL) limited the exchange of O_2_ between the biofilm and the surrounding water (Fig. 1 and 4). In dark, O_2_ was depleted within the upper 1.5 mm and the areal dark respiration rate was calculated to 0.039 nmol O_2_ cm^−2^ s^−1^. The fluxes of O_2_ increased with irradiance until saturation was reached at a downwelling photon irradiance of ~100 µmol photons m^−2^ s^−1^, where the top of the biofilm experienced O_2_ concentrations >450% of air saturation (Fig. 4). The O_2_ concentration profiles for diffuse and collimated light were very similar, although O_2_ penetrated deeper under diffuse light, especially at the highest photon irradiances (500 and 1000 µmol photons m^−2^ s^−1^) (Fig. 4). The coral sediment had a ~1-1.4 mm thick DBL; dark respiration was similar to the biofilm (0.037 nmol O_2_ cm^−2^ s^−1^), while saturation of photosynthesis was reached at a higher downwelling photon irradiance of ~200 µmol photons m^−2^ s^−1^ (Fig. 3). The more variable DBL thickness in the coral sediment varied independently of irradiance and was most likely a result of the very heterogeneous surface topography (Fig. 4). At incident irradiances >200 µmol photons m^−2^ s^−1^ the O_2_ productive zone was stratified under both diffuse and collimated light, with an O_2_ concentration maximum of ~600% air saturation ~1.7 mm below the sediment surface (Fig. 4). Photosynthesis was apparently distributed in two major layers, a ~0.5 mm thick layer at the sediment surface, and a ~1 mm thick layer peaking 2 mm below the sediment surface (Fig. 4 and S3). The O_2_ concentration profiles for diffuse and collimated light were similar at low to moderate irradiance, then showed a deeper O_2_ penetration depth under diffuse light at incident irradiance >500 µmol photons m^−2^ s^−1^ in comparison to O_2_ profiles measured under collimated light (Fig. 4). The O_2_ profiles in the coral sediment showed relatively high standard deviations, possibly due to a more patchy distribution of the photosynthetic organisms within the sediment and overall variability in the sediment grain size and surface topography.

**Figure 4.**
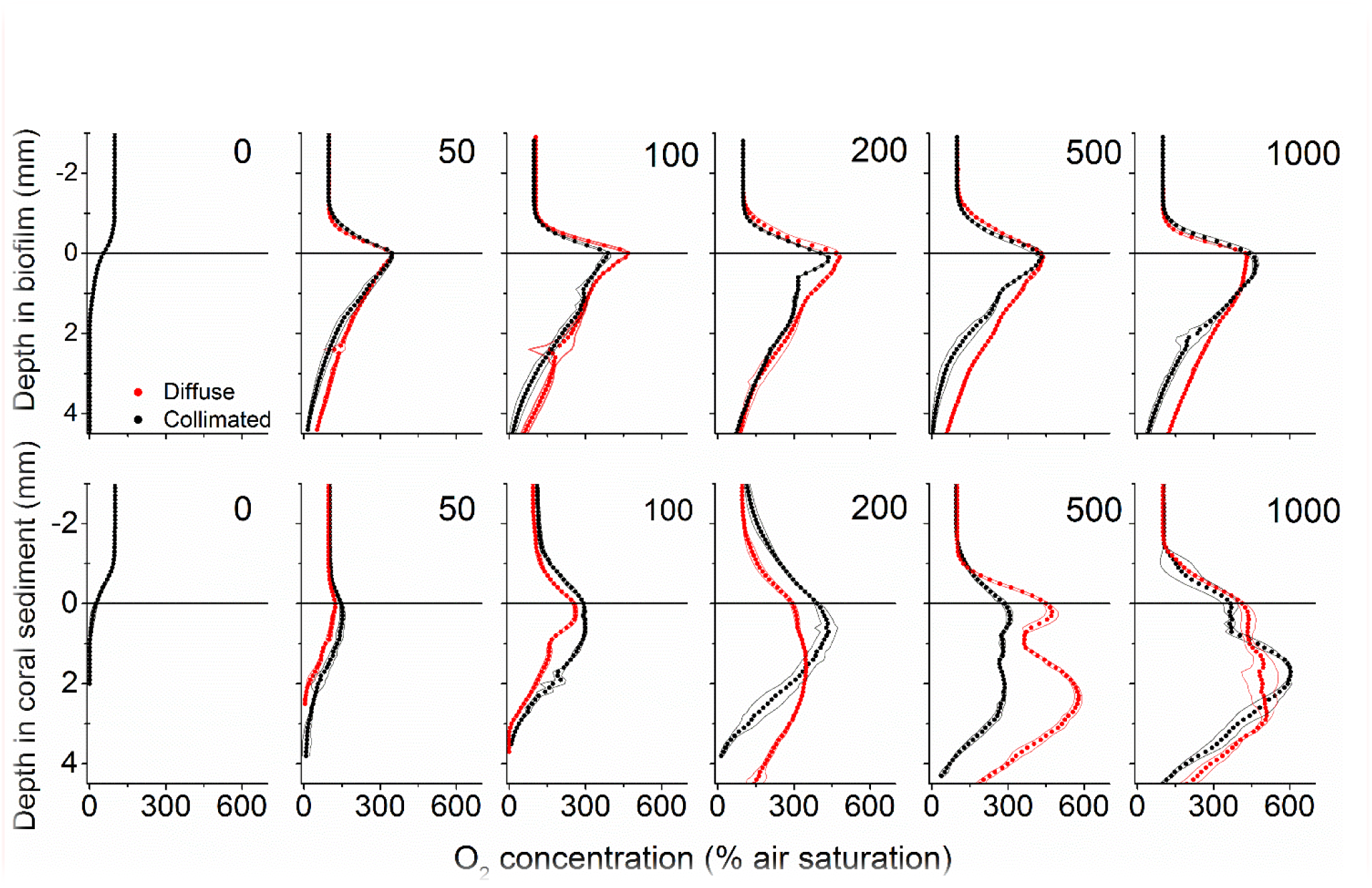
**Vertical microprofiles of O_2_ concentration** In biofilm (upper panel) and coral sediment (lower panel). Red and black symbols represent measurements under diffuse and collimated light, respectively, while numbers in panels denote downwelling photon irradiance in µmol photons m^−2^ s^−1^. The line in y=0 indicates the biofilm/sediment surface. Symbols represent mean values, while dashed lines represent ± 1 S.D. (*n*=3).

In both biofilm and coral sediment, the surface temperature increased relative to the overlaying seawater with increasing irradiance. The local heating was dissipated by heat transfer over a ~3 mm thick thermal boundary layer (TBL) into the overlaying seawater and into deeper sediment layers (Fig. 5 and 6). Robust measurements of biofilm/sediment heating could only be obtained at incident photon irradiances of ≥200 µmol photons m^−2^ s^−1^ (≥500 µmol photons m^−2^ s^−1^ for the coral sediment 397 under collimated light). At the highest irradiance (1000 µmol photons m^−2^ s^−1^), the biofilm surface 398 was 0.51°C and 0.41°C warmer than the overlaying water, while the coral sediment surface was 0.53°C and 0.48°C warmer than the surrounding water for diffuse and collimated light, respectively. Similar temperature profiles were observed between collimated and diffuse light, although a slightly enhanced surface heating and thus a higher efflux of heat was observed under diffuse light (Figure 5). Comparing the slope of the surface warming vs. vector irradiance under diffuse and collimated light, respectively, diffuse light had a greater impact on surface warming by 30% and 27% in the biofilm and in the coral sediment, respectively (Fig. 6).

**Figure 5.**
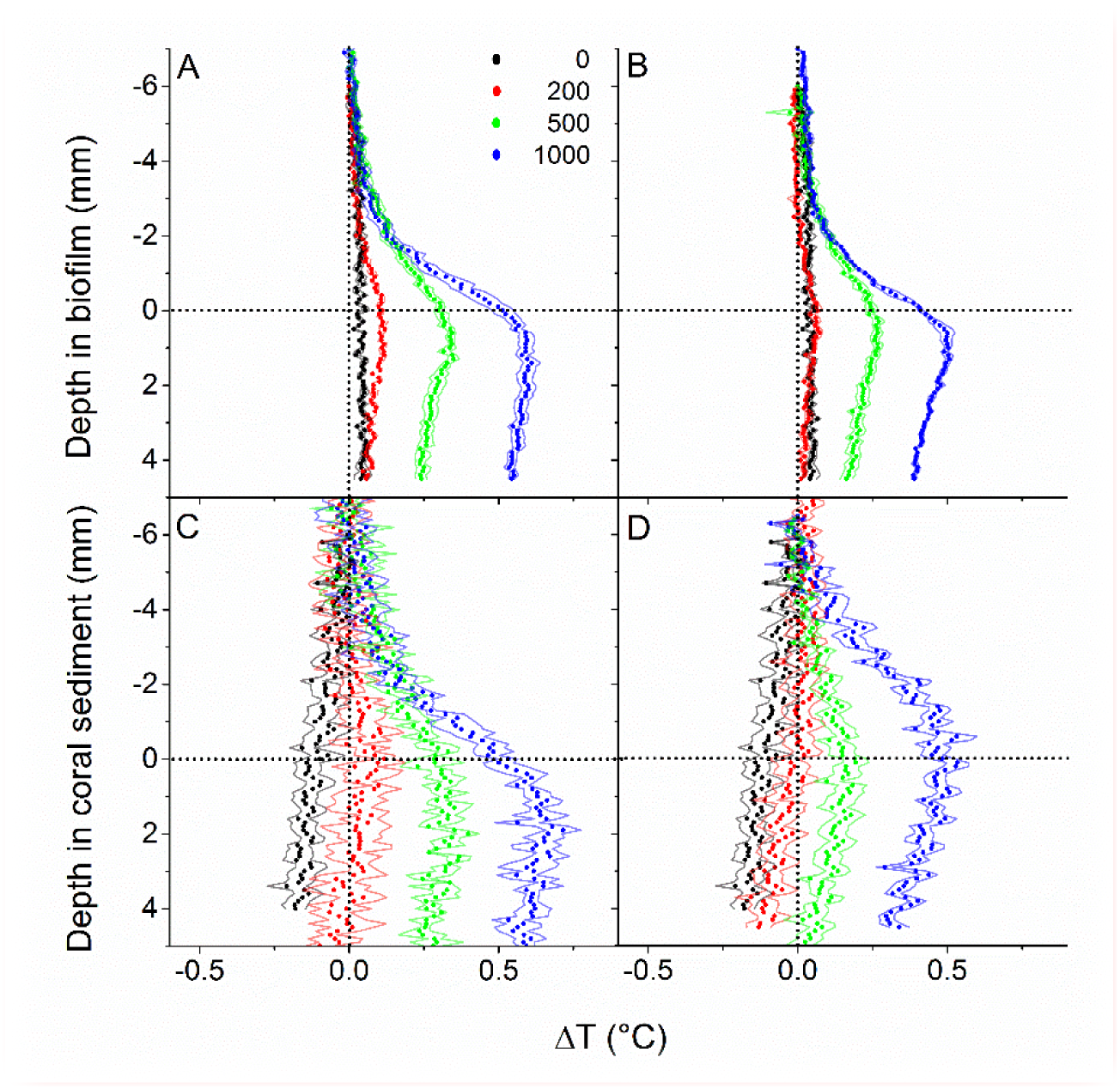
**Vertical depth profiles of temperature change,** ΔT (in °C) measured in biofilm (upper panels) and coral sediment (lower panel) at downwelling photon irradiances of 0, 200, 500 and 1000 µmol photons m^−2^ s^−1^ under collimated (A, C) and diffuse light (B, D). Symbols represent means, while dashed lines indicate ± 1 S. D. (*n*=3). The dotted line in y=0 indicates the sediment surface, while the dotted line in x=0 indicates a 0°C temperature change.

**Figure 6.**
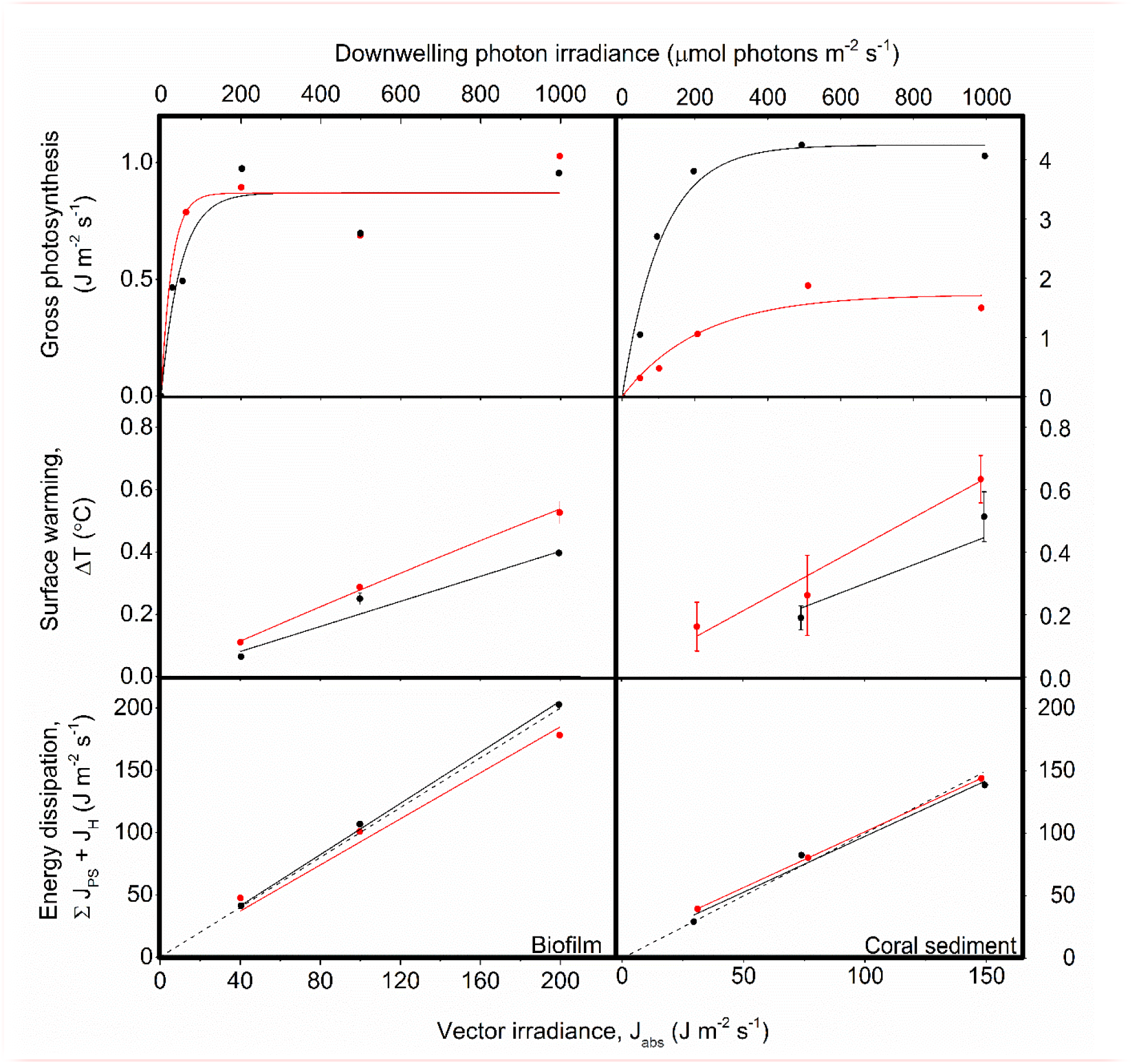
**Energy conversion by photosynthesis, heat dissipation and the sum of photosynthesis and heat dissipation versus downwelling irradiance** In biofilm (left panels) and corals sediment (right panels). Red symbols and lines show data for diffuse illumination, while black symbols and lines show data for collimated illumination. ***A, B***) Areal gross photosynthesis rates (in J m^−2^ s^−1^) measured at downwelling photon irradiances of 0, 50, 100, 200, 500 and 1000 µmol photons m^−2^ s^−1^, and then fitted with a saturated exponential model ((Webb *et al*., 1974)); CS: R^2^_diff_ = 0.92, R^2^_coll_ = 0.97; BF: R^2^ = 0.88 for both diffuse and collimated; n = 3). ***C, D***) Temperature gradients (in °C) between the ambient seawater and the sediment surface (flow = 0.3-0.4 cm s-1), measured at vector irradiances of 30, 75 and 149 J m^−2^ s^−1^ or 40, 100 and 200 J m^−2^ s^−1^ for the coral sediment and biofilm, respectively. Data points show means ± SD (*n* = 3); CS: R^2^_diff_ = 0.99, R^2^_coll_ = 0.96; BF: R2 = 0.99 for both diffuse and collimated light. ***E, F***) The summed energy dissipation of the system (in J m^−2^ s^−1^), i.e., the sum of energy conserved by photosynthesis and energy dissipated as heat, measured at vector irradiances of 30, 75 and 149 J m^−2^ s^−1^ and 40, 100 and 200 J m^−2^ s^−1^ for the coral sediment and biofilm, respectively. The dashed line represents a 1:1 relationship between the incoming and outgoing energy of the system (i.e. the theoretically expected relationship). CS: R^2^_diff_ = 0.99, R^2^_coll_ = 0.96; BF: R^2^ = 0.99 for both diffuse and collimated light; (*n* = 3).

### Photosynthesis

Maximal volume-specific gross photosynthesis rates of the biofilm ranged between 7.0 nmol O_2_ cm^−3^ s^−1^ and 8.7 nmol O_2_ cm^−3^ s^−1^ (collimated and diffuse light, respectively) under low irradiance (50-200 µmol photons m^−2^ s^−1^), while rates decreased at photon irradiances of >200 µmol photons m^−2^ s^−1^ (Fig. S3A). The thickness of the photic zone generally increased with increasing photon irradiance and varied from 0.4-1.2 mm in the biofilm under diffuse light and from 0.2-0.9 mm under collimated light.

In the coral sediment, the highest volume-specific rates of photosynthesis were measured within the upper 1 mm, with maximal gross photosynthesis rates of 11.97 nmol O_2_ cm^−3^ s^−1^ at the sediment surface under collimated light and 3.05 nmol O_2_ cm^−3^ s^−1^ at a depth of 0.6 mm under diffuse light (Fig. S3B). The photic zone in the coral sediment increased with increasing irradiance and ranged in thickness from 1.5 to 3 mm under diffuse light and from 2 to 3.5 mm under collimated light. The apparent stratification in O_2_ concentration found in the coral sediment, was confirmed in the profiles of gross photosynthesis, with peaks in gross photosynthesis in the upper 1 mm, and 1.5-2.5 mm from the surface at photon irradiances >50 µmol photons m^−2^ s^−1^ (Fig. S3B).

Under low photon irradiance <200 µmol photons m^−2^ s^−1^ in the biofilm, the area specific gross photosynthesis rate (PG) was higher under diffusive illumination, while PG under diffuse and collimated illumination were similar at higher irradiances (Fig. 6). In contrast, PG in the coral sediment was generally in the range of 3-4 times lower under diffuse-compared to collimated light (Fig. S3B; Fig. 6B). We note that the gross photosynthesis measurements in the coral sediment under diffuse light were performed at the University of Technology Sydney (UTS) rather than on Heron Island Research Station (HIRS), where the rest of the measurements took place. This apparently resulted in a changed microbial community structure of the coral sediment. We speculate that the transport from Heron Island created prolonged anoxic conditions throughout the sediment and this might have caused this possible change in community structure. These measurements were therefore excluded when calculating the light energy budget for diffuse light in the coral sediment.

### Energy budgets

The photosynthesis-irradiance (PE) curve of the coral sediment measured in diffuse light increased with increasing light intensity with an initial slope of 0.05±0.01, until reaching an asymptotic saturation level at *J_PS, max_* = 1.72±0.20 J m^−2^ s^−1^ at a downwelling photon irradiance of ~300 µmol photons m^−2^ s^−1^ (Fig. 6B). In contrast, the PE-curve of the coral sediment in collimated light increased with the with a slope of 0.26±0.04, reaching a maximum saturation value of *J_PS, max_* = 4.24±0.23 J m^−2^ s^−1^, at downwelling photon irradiance ~110 µmol photons m^−2^ s^−1^ (Fig. 6B). In the biofilm, the onset of photosynthesis saturation occurred already at a downwelling photon irradiance of ~50 µmol photons m^−2^ s^−1^, where *J_PS, max_* reached an asymptotic saturation level of 0.87 J m^−2^ s^−1^ for both diffuse and collimated light (Fig. 6A).

Sediment surface warming increased linearly with irradiance under both diffuse and collimated light with average slopes of CSα_diff_ = 4.33·10^−3^ °C (J m^−2^ s^−1^)^−1^ and CSα_coll_ = 2.14·10^−3^ °C (J m^−2^ s^−1^)^−1^ in the coral sediment, as compared to BFα_diff_ = 2.77·10^−3^ °C (J m^−2^ s^−1^)^−1^ and BFα_coll_ = 2.0·10^−3^ °C (J m^−2^ s^−1^)^−1^ in the biofilm (Fig. 6C,D). Surface warming was stronger under diffuse light as compared to collimated light in both sediments (Fig. 5, Fig. 6C,D).

The summed flux of energy conserved by photosynthesis and dissipated as heat (J_PS_ + J_H_) serves as a control to determine the potential deviations between absorbed and dissipated energy (Fig. 6E, F). Dissipation of energy from the system increased linearly with increasing vector irradiance with slopes in the coral sediment of 0.89±0.003 and 0.89±0.120, for diffuse and collimated light respectively, and slopes in the biofilm of 0.93 and 1.03, for diffuse and collimated light respectively. Thus, the method is very close to the theoretical expected slope of 1 (dashed line), where the outgoing/used energy equals the incoming light energy.

About 29% of the incident light energy was back-scattered from the coral sediment surface and thus not absorbed, whereas the surface reflection was only ~2% of the incident irradiance in the biofilm (Fig. 7; Fig. S2). The fraction of energy conserved by photosynthesis decreased with increasing irradiance in both biofilm and sediment (Fig. 7; 8). Over the investigated incident irradiances (200 – 1000 µmol photons m^−2^ s^−1^), photosynthetic energy conservation in the coral sediment illuminated with diffuse light decreased from 6.7% to 2.0% of the incident light energy, favouring heat dissipation (which increased from 63.1% to 67.8%), and from 9.3% to 2.1% of the incident light energy under collimated light (where heat dissipation increased from 62.6% to 69.8%) (Fig. 7; Table S2).

**Figure 7.**
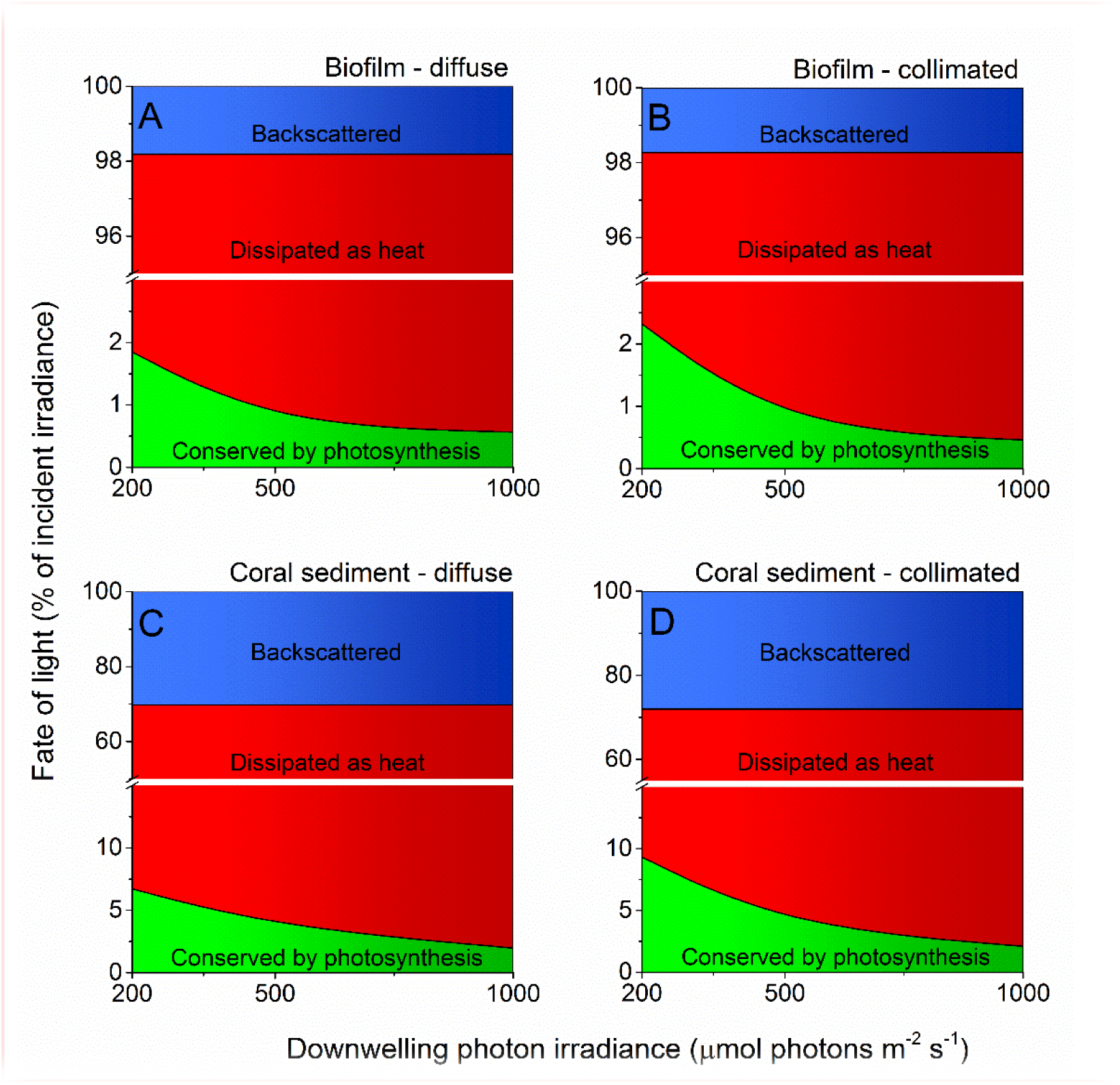
**Light energy budgets** for biofilm (A, B) and coral sediment (C, D) in percent of the incident light energy calculated at downwelling photon irradiance (PAR) of 200, 500 and 1000 µmol photons m^−2^ s^−1^, under diffuse (A, C) and collimated (B, D) incident light. The amount of light backscattered from the sediment surface is shown in blue, while the amount of light energy dissipated as heat and via photosynthesis is shown in red and green, respectively. Notice the break on the y-axis. We assumed similar GPP under diffuse and collimated light in the calculations for the coral sediment under diffuse light (*see Table S1 and Figure S3*).

The proportion of incident light energy that was conserved via photosynthesis, was much lower in the biofilm, where 1.9% and 2.3% (diffuse and collimated light, respectively) of the incident light energy was conserved, whereas 96.3% and 96.0% of the incident light energy was dissipated as heat, respectively (Fig. 7; Table S1). At an incident irradiance of 1000 µmol photons m^−2^ s^−1^, only 0.6% and 0.5% of the incident energy was conserved by photosynthesis, while 97.6% and 97.8% was dissipated as heat under diffuse and collimated light, respectively (Fig. 7; Table S2).

The maximum photochemical energy conservation in the coral sediment was observed at an incident irradiance of ~100 µmol photons m^−2^ s^−1^ (18.1% of the absorbed light energy), whereas the biofilm had maximum energy conservation through photosynthesis (14.7% of the absorbed light energy) at the lowest measured incident irradiance (50 µmol photons m-2 s-1) (Fig. 8). In addition, the biofilm had higher photosynthetic efficiencies under diffuse light compared to collimated light at low light intensities (<200 µmol photons m^−2^ s^−1^) (Fig. 8).

**Figure 8.**
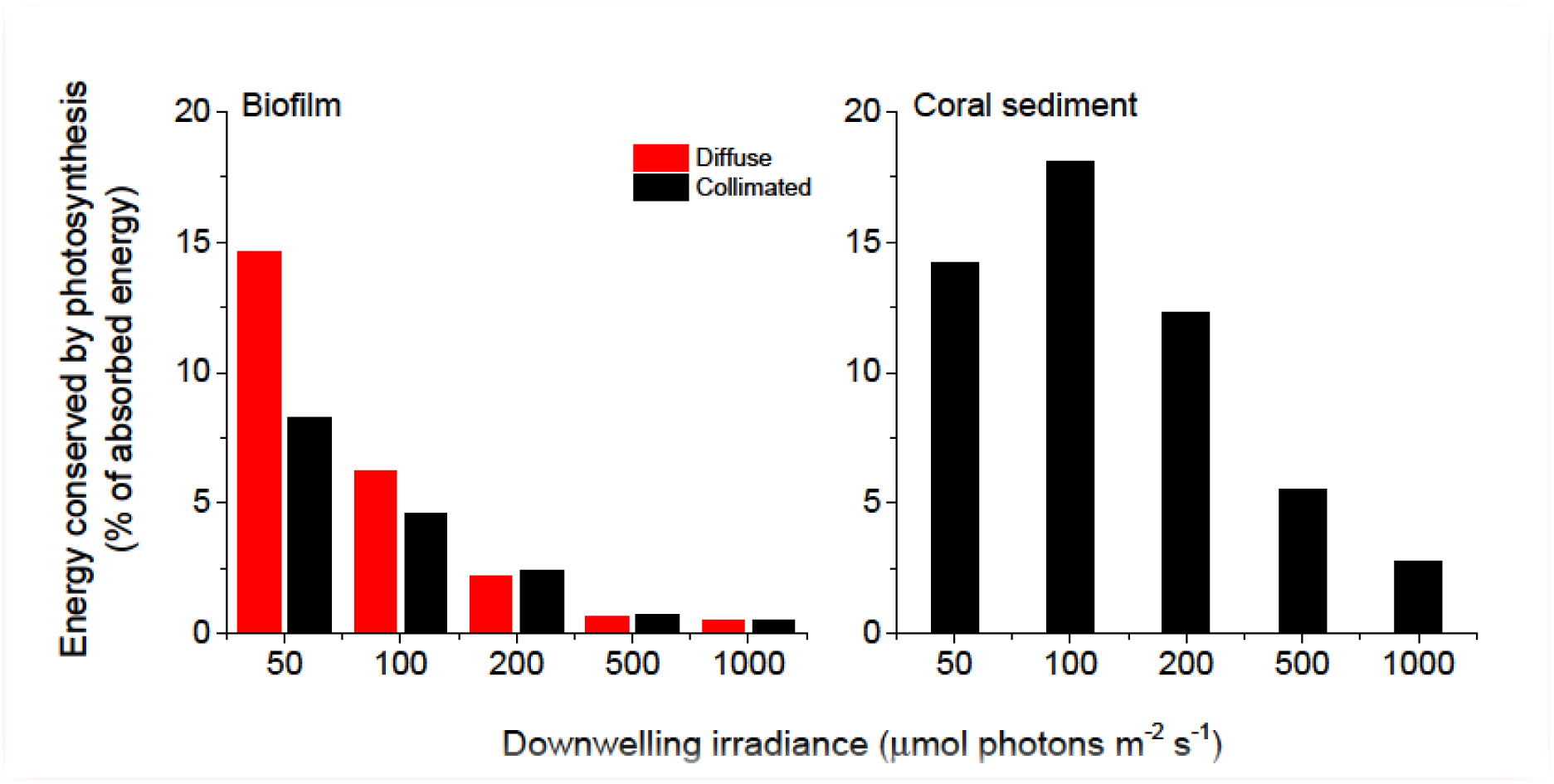
**Photosynthetic energy conservation efficiencies** (in % of the absorbed light energy) of the biofilm (*left panel*) and the coral sediment (*right panel*), measured at incident photon irradiances of 50, 100, 200, 500 and 1000 µmol photons m^−2^ s^−1^, under diffuse (red bars) and collimated (black bars) light.

The photosynthetic efficiencies of biofilm and coral sediment under light-limiting conditions (J_ABS_→0), ɛPS,_max_, were calculated from the initial slope of the areal PG vs. vector irradiance curve to 26.2% of the absorbed light energy (CS, collimated light) compared to 16% and 9.0% of the absorbed light energy (BF, diffuse and collimated light, respectively).

## Discussion

We present a closed radiative energy budget of a heterogeneous coral reef sediment and compare it to the radiative energy budget of a flat dense biofilm (Fig. 6 and S4). The closed light energy budgets were measured under both diffuse and collimated illumination to test potential effects of the directionality of light on the photosynthetic efficiencies of the phototrophs. We found that a higher fraction of the absorbed light energy was conserved by photosynthesis in the heterogenous loosely organized coral sediment, while the radiative energy budgets of both sediment types were highly dominated by dissipation of heat.

### Light

The thin highly pigmented cyanobacterial biofilm was growing on the surface of a fine-grained (125-250 µm) dark sandy sediment, whereas the photosynthetic microorganisms exhibited a more patchy distribution within the large-grained (100-500 µm) bright and highly scattering coral sediment. This structural difference between the two systems led to a ~15 times higher surface reflection and a decreased energy absorption in the coral sediment compared to the biofilm that displayed >8 times higher light attenuation coefficients. As previously shown (Lassen *et al*., 1992b; Kühl and Fenchel, 2000) the scalar irradiance at, or immediately below, the surface increased, and the spectral composition was altered relative to the incident irradiance (Fig. 2 and 3). Such increase in scalar irradiance in the near surface layer is due to intense multiple scattering by particles (biotic and abiotic) causing a local photon path-length increase and thus a prolonged residence time of scattered photons in the surface layers that also receive a continuous supply of incident photons from the light source (Kühl and Jørgensen, 1994). This effect can be further enhanced in the presence of exopolymers with a slightly higher refractive index than the surrounding seawater leading to photon trapping effects (Kühl and Jørgensen, 1994; Decho *et al*., 2003). Furthermore, the structural difference between the loosely organized CaCO3 particles compared to the flat biofilm could possibly result in differences in the reflection characteristics from the uppermost layers. In the biofilm, the flat homogeneous surface reflects light relatively uniformly, with some ratio between specular vs. diffuse reflection. However, in the heterogeneous coral sediment a higher degree of forward scattering will most likely be present as the angle of reflection will be more complex due to the roughness of the surface, resulting in a deeper penetration of light in the coral sediment.

### Temperature

We directly measured both the upward and downward heat dissipation of radiative energy (Fig. 5). Previous studies of energy budgets ignored the downward heat flux (Al-Najjar *et al*., 2010; Al-Najjar *et al*., 2012), and although Jimenez *et al*. (2008) estimated the downward heat dissipation from a theoretical model considering the physical properties of heat transfer in coral skeleton, this is to the best of our knowledge the first study of energy budgets of phototrophic systems, where the complete heat balance was directly measured. Over a range of incident irradiances, the downward heat flux was the same order of magnitude as the upward heat flux in both biofilm and coral sediment and thus is an important parameter when compiling light energy budgets for the photic zone in benthic systems (Fig. 5).

The majority of the absorbed light energy was dissipated as heat (Fig. 7; Table S1) with a linear relationship between increasing incident irradiance and heat dissipation under both diffuse and collimated light, albeit with a ~30% enhanced surface warming under diffuse light as compared to collimated light (Fig. 5 and 6). Apparently, diffuse light was absorbed more efficiently in the uppermost layers, increasing the local photon density and residence time in these layers resulting in increased energy deposition and surface temperatures. This was supported by a higher heat flux into the water column under diffuse light, and a higher heat flux into the sediment under collimated light (data not shown). At increasing irradiances the surface temperature of the sediments exceeded the surrounding water temperature and convective heat transport occurred over the TBL (Fig. 5) (Jimenez *et al*., 2011). While we cannot dissect the observed heat dissipation into particular mechanisms and their relative magnitude, part of such dissipation in optically dense biofilms and sediments involves non-photochemical quenching (NPQ) processes that protect the photosynthetic apparatus under high irradiance by channelling excess light energy into heat dissipation (Falkowski and Raven, 2007; Al-Najjar *et al*., 2012). The heat fluxes from the photic zone were generally higher in the biofilm when compared to the coral sediment, due to the lower reflection and thus higher absorption in the biofilm Fig. S4). However, when normalizing the heat fluxes to the absorbed light energy (which was 33% higher in the biofilm than in the coral sediment) the heat dissipation was of similar magnitude, and variations in heat flux values between the sediment and biofilm became <15%. The degree of heat dissipation therefore seems tightly correlated to the quantity of absorbed energy.

### Photosynthesis

The overall photosynthetic efficiency of the biofilm and coral sediment decreased with increasing incident irradiance, with photosynthesis exhibiting saturation at higher irradiance under both diffuse and collimated light (Fig. 6). The highest energy storage efficiency of the coral sediment was observed under light-limiting conditions (<200 µmol photons m^−2^ s^−1^) (Fig. 7, 8), and the coral sediment generally exhibited high light use efficiencies that were comparable to those observed in corals at equivalent incident photon irradiances (Brodersen *et al*., 2014). The photosynthetic activity in the coral sediment was stratified at incident irradiances >50 µmol photons m^−2^ s^−1^ under both diffuse and collimated light (Fig. 4). This stratification could be a result of different factors influencing the photosynthetic activity such as steep light attenuation over depth, locally high volume-specific rates of metabolic activity, higher local biomass of phototrophs and diffusion limitation of metabolic products and substrates (Kühl *et al*., 1996; Kühl and Fenchel, 2000; Al-Najjar *et al*., 2012). Such vertical stratification has also been associated with phototactic responses to light (Whale and Walsby, 1984; Lassen *et al*., 1992b), where motile photosynthetic organisms migrate to an optimal depth for photosynthesis at a given irradiance, where the available light is neither limiting nor inhibiting the rate of photosynthesis (Al-Najjar *et al*., 2012). These migration patterns are well documented both as photo-and aero-tactic responses and to escape from e.g. toxic levels of sulphide (Kühl *et al*., 1994; Bebout and Garcia-Pichel, 1995). The two photosynthetic active layers were situated at the sediment surface and ~2 mm below (~0.5 mm and 1 mm thick layers, respectively; Fig. 3).

The area-specific rates of gross photosynthesis of the coral sediment were ~4 times higher than in the biofilm, due to a ~3 times deeper euphotic zone and slightly higher volume-specific photosynthesis rates in the coral sediment than in the biofilm (Fig. 6, 7 and Fig. S3). Consequently, the coral sediment reached an asymptotic maximum in PG in terms of energy dissipation via photosynthesis of ~4.2 J m^−2^ s^−1^ as compared to only ~0.9 J m^−2^ s^−1^ in the biofilm (Fig. 6). The E_k_ value, i.e., the irradiance at the onset of photosynthesis saturation, was >2 higher in the coral sediment compared to the Danish biofilm, which reflects the different *in situ* light conditions experienced by the two systems in their respective geographic locations (Denmark: 55°N, Heron Island: 23°S). Thus, the dense biofilm appeared acclimated to low irradiances as previously shown (Kühl *et al*., 1996; Kühl and Fenchel, 2000; Al-Najjar *et al*., 2012) where highly reduced quantum efficiencies are seen at increasing irradiance due to the employment of e.g. NPQ processes. Accordingly, the coral sediment maintained higher photosynthetic efficiencies, even at high irradiance. This could in part be explained by the high scattering in the sediment particles that creates a more even spread of the light field over the sediment matrix and a more dispersed photic zone; a factor that have been speculated to be responsible for the high photosynthesis in coral tissues (Brodersen *et al*., 2014; Wangpraseurt *et al*., 2014a). A more homogenous distribution of light would create a larger region where light is neither limiting nor inhibiting photosynthesis. Thus, the loosely organized more heterogenous coral sediment apparently 579 exhibit a more open, canopy-like organization compared to the dense biofilm.

Community photosynthesis is generally higher than that of individual phytoelements (Binzer and Sand-Jensen, 2002; Binzer and Middelboe, 2005; Binzer *et al*., 2006) and in addition, higher community photosynthesis have been found under diffuse illumination in open canopy systems which was explained by a more even light field inside the canopy (Gu *et al*., 2002; Brodersen *et al*., 2008). In spite of this difference in overall acclimation to light, a decrease in the surface layer photosynthesis was seen in the coral sediment at an incident irradiance of 500 µmol photons m^−2^ s^−1^, which could either reflect the heterogeneity and patchiness of the phototrophs found in the sediment, or could point to a possible migration of motile phototrophic organisms. Migration as a phototactic response is recognized as an effective mechanism for regulating photon absorption across different systems such as terrestrial plants (Wada *et al*., 2003) and microphytobenthic assemblages (Serodio *et al*., 2006; Cartaxana *et al*., 2016a; Cartaxana *et al*., 2016b), and similar phototactic response have been shown in coral tissues where the *in hospite* light environment can be modulated by host tissue movement (Wangpraseurt *et al*., 2014a; Wangpraseurt *et al*., 2016). Downward migration at high irradiances is probably correlated with increasing photic stress e.g. by the formation of reactive oxygen species that can affect photosystem II by damaging (or preventing the repair of) the D1 protein in these layers (Hihara *et al*., 2001; Nishiyama *et al*., 2001; Aarti *et al*., 2007; Latifi *et al*., 2009; Al-Najjar *et al*., 2010). Several ways to counter such photic stress exists. One of the most effective short-term responses to photic stress is to employ non-photochemical quenching (NPQ) in which photons are emitted as heat when cells experience over-saturating photon fluxes. Another strategy to avoid photodamage is to upregulate the expression of sun-protective pigments such as β-carotenes (Zhu *et al*., 2010), which were found in significant amounts by HPLC analysis of the coral sediment (Fig. S1).

Photosynthetic energy conservation was higher under collimated light as compared to illumination with diffuse light at moderate irradiance (200 µmol photons m^−2^ s^−1^) (Fig. 6). This finding correlates with previous studies of individual terrestrial leaves reporting 10-15% higher energy conservation via photosynthesis under collimated-relative to diffuse light (Brodersen *et al*., 2008) and in corals it has been shown that gross photosynthesis was 2-fold higher under direct vs. diffuse light (Wangpraseurt and Kühl, 2014). In terrestrial leaves, the more efficient utilization of collimated light has been ascribed to specialized tissue structures such as columnar palisade cells (Vogelmann and Martin, 1993), that increase the absorptance of direct light over diffuse light (Brodersen and Vogelmann, 2007). Furthermore, light-induced chloroplast movement has been shown to be less effective under diffuse illumination (Gorton *et al*., 1999; Williams *et al*., 2003). In corals the higher photosynthesis at the tissue surface was explained by optical properties enhancing the scalar irradiance near the surface under direct illumination (Wangpraseurt and Kühl, 2014). This tendency changed dramatically in the dense photosynthetic biofilm at light-limiting conditions (⩽100 µmol photons m^−2^ s^−1^) favouring effective light utilization under diffuse light (Fig. 7). Thus, the optical properties and the structural organization of phytoelements seem tightly linked to the photosynthetic quantum efficiencies across different systems and light angularity may therefore elicit differential photosynthetic responses depending on system and on scale.

### Conclusion

Our results showed that a higher fraction of the absorbed light energy was conserved by photosynthesis in the heterogeneous coral sediment due to a deeper photic zone and slower saturation of photosynthesis with increasing irradiance as compared to the flat and highly absorbing biofilm. The balanced radiative energy budget of biofilm and coral sediment was highly dominated by heat dissipation and the efficiency of photosynthetic energy conservation decreased with increasing irradiance. Several variances were found between illumination with diffuse or collimated light: i) diffuse light enhanced dissipation of heat (~30%) in the upper sediment layers as compared to collimated light; ii) at low incident irradiance (200 µmol photons m^−2^ s^−1^) photosynthetic energy conservation was higher (3-4% of the absorbed light energy) in collimated light as compared to diffuse light; a tendency that dramatically changed in the photosynthetic biofilm at low and light-limiting incident irradiances (≤100 µmol photons m^−2^ s^−1^) favouring effective light utilization under diffuse light (up to a ~2-fold increase). However, cyanobacterial and diatom dominated mats have been shown to migrate vertically employing photo-and/or chemo-tactic responses (Richardson and Castenholz, 1987; Bhaya, 2004; Serodio *et al*., 2006; Coelho *et al*., 2011; Cartaxana *et al*., 2016a) and the motility of the phototrophs was not considered here. Thus, there is a need to further explore how vertical migration affects the radiative energy balance and thereby the light use efficiency in microbenthic systems such as sediments and biofilms.

**Definition of abbreviations and parameters**

**Definition of abbreviations and parameters**
| **Abbreviation** | **Definition** | **Unit** |
| --- | --- | --- |
| **DBL** | Diffusive boundary layer |  |
| **TBL** | Thermal boundary layer |  |
| **PAR** | Photosynthetically active radiation (420-700nm) |  |
| **BF** | Biofilm from Limfjorden, Denmark |  |
| **CS** | Coral sediment from Heron Island lagoon, Australia |  |
| **J<sub>NPP↑</sub>** | Upward O<sub>2</sub> flux from photic zone | nmol O<sub>2</sub> m<sup>-2</sup> s<sup>-1</sup> |
| **J<sub>NPP↓</sub>** | Downward O<sub>2</sub> flux from photic zone | nmol O<sub>2</sub> m<sup>-2</sup> s<sup>-1</sup> |
| **J<sub>NPP</sub>** | Total net photosynthesis in photic zone<br>(J<sub>NPP↑</sub> - J<sub>NPP↓</sub>) | nmol O<sub>2</sub> m<sup>-2</sup> s<sup>-1</sup> |
| **φ** | Sediment porosity | Dimensionless |
| **P<sub>G</sub>(z)** | Volume-specific rate of gross photosynthesis | nmol O<sub>2</sub> cm<sup>-3</sup> s<sup>-1</sup> |
| **P<sub>G</sub>** | Area-specific rate of gross photosynthesis | nmol O<sub>2</sub> cm<sup>-2</sup> s<sup>-1</sup> |
| **J<sub>PS</sub>** | Area-specific rate of gross photosynthesis in energy terms | J m<sup>-2</sup> s<sup>-1</sup> |
| **J<sub>H↑</sub>** | Upward heat flux from photic zone | J m<sup>-2</sup> s<sup>-1</sup> |
| **J<sub>H↓</sub>** | Downward heat flux from photic zone | J m<sup>-2</sup> s<sup>-1</sup> |
| **J<sub>H</sub>** | Total heat flux out of photic zone J<sub>H↑</sub> + J<sub>H↓</sub> | J m<sup>-2</sup> s<sup>-1</sup> |
| **E<sub>d</sub>(λ)** | Spectral downwelling irradiance | Counts nm<sup>-1</sup> |
| **E<sub>0</sub>(λ)** | Spectral scalar irradiance | % of E<sub>d</sub>(λ) |
| **J<sub>ABS</sub>** | Absorbed light energy | J m<sup>-2</sup> s<sup>-1</sup> |
| **R** | Reflection | % |
| **K<sub>0</sub>(PAR)** | Diffuse attenuation coefficient of PAR scalar irradiance | mm<sup>-1</sup> |
| **K<sub>0</sub>(λ)** | Spectral attenuation coefficient of scalar irradiance | mm<sup>-1</sup> |
| **E<sub>k</sub>** | Photochemical light acclimation index, i.e., the irradiance at onset of photosynthesis saturation | μmol photons m<sup>-2</sup> s<sup>-1</sup> |

## Acknowledgements

We thank P. J. Ralph, P. Brooks, M. Zbinden and other colleagues at University of Technology Sydney (C3, UTS) for access to laboratory facilities, technical support and help with HPLC analysis of the coral sediment. We thank the staff at Heron Island Research Station for technical assistance during the field work. V. Schrameyer, D. Wangpraseurt and D. A. Nielsen are thanked for thoughtful discussions. The research was conducted under research permits for field work on the Great Barrier Reef, Australia (G11/34670.1 and G09/31733.1) and was funded by the Danish Council for Independent Research | Natural Sciences (MK), the *Knud Højgaards Fund, the Oticon Foundation, Thorsons Travel Grant and Københavns Universitets Fælleslegat* (ML, KEB).

### Author contributions

ML, KB and MK designed the experiments; ML and KB performed experiments; ML, KB and MK analyzed data; ML wrote the paper with editorial inputs from KB and MK.

### Supplementary material

**Figure S1.** Depth distribution of major photopigments.

**Figure S2.** PAR surface reflectance.

**Figure S3.** Depth profiles of volumetric gross photosynthesis rates.

**Figure S4.** Absorbed light energy vs. downwelling irradiance.

**Table S1.** Calculated fluxes of O_2_, heat and absorbed light energy.

**Table S2.** Calculated fractions of incident light reflected, conserved in photosynthesis or dissipated as heat.

